# Sex-specific DNA methylation in adult skeletal muscle

**DOI:** 10.64898/2026.03.16.712025

**Authors:** Clara Martínez Mir, Ruben Boers, Joost Gribnau, Anna Alemany, Fanny Sage, Niels Geijsen

**Affiliations:** Department of Anatomy and Embryology, Leiden University Medical Center, 2333 Leiden, The Netherlands; The Novo Nordisk Foundation Center for Stem Cell Medicine (reNEW), Leiden University Medical Center, The Netherlands; Department of Developmental Biology, Erasmus University Medical Center Rotterdam, Rotterdam, The Netherlands

## Abstract

DNA methylation is a key epigenetic mechanism influencing gene regulation and cellular identity. In skeletal muscle, methylation contributes to fiber-type specification, metabolic programming, and satellite cell function, with evidence of sex-specific differences. Here, we investigated whether spatial regionalization of gene expression along the proximal-distal axis of the tibialis anterior (TA) is mirrored by corresponding patterns of DNA methylation. Using MeDseq on TA sections from muscles previously analyzed by spatial transcriptomics, we profiled methylation across transcriptional start sites (TSS), gene bodies, and regulatory elements. Despite robust spatial differences in transcriptomes, methylation patterns were largely uniform along the proximal-distal axis, indicating that DNA methylation does not underlie regional gene expression in adult TA muscle. In contrast, sex emerged as the primary determinant of methylation variation. Male muscles exhibited widespread hypermethylation at TSS, gene-bodies and regulatory regions, corresponding with sex-specific transcriptional programs, including glycolytic fiber enrichment in males and oxidative fiber markers in females. Notably, chromatin- and methylation-associated regulators such as *Setd7, Gsk3a*, and *Bmyc* were upregulated in males, suggesting mechanisms linking transcriptional control to epigenetic state. These findings highlight that while spatial gene expression is transcriptionally driven, sex-specific epigenetic programs dominate adult skeletal muscle, underscoring the need to consider sex in multi-omic studies of muscle biology.

## Introduction

DNA methylation is an epigenetic modification that shapes transcriptional outputs and cellular identity. By modulating chromatin accessibility and transcription factor binding, methylation contributes to both the activation and repression of gene expression, depending on genomic context^1^. Classically, CpG methylation at promoter regions and transcription start sites (TSSs) is associated with transcriptional repression, whereas gene body methylation frequently correlates with active transcription and may prevent spurious initiation within coding region^2,3^. Beyond promoters and gene bodies, methylation dynamics at distal regulatory elements, including enhancers, further refine gene expression programs and enable cell-type-specific regulatory landscapes.

In skeletal muscle, DNA methylation plays essential roles in maintaining tissue homeostasis and supporting the functional diversity of myofibers. Satellite cells require *Dnmt1*-dependent maintenance methylation to preserve quiescence and enable efficient regeneration^4^. At the myofiber level, distinct methylation patterns have been observed between slow- and fast-twitch fiber types^5^, and these profiles can be remodeled by external stimuli such as exercise training that induce fiber switching between fast and slow-twitch^6^. In addition, several recent studies have reported pronounced sex differences in the skeletal muscle methylome, linking differential methylation to variation in muscle function and substrate metabolism between males and females^7-9^.

Building on the transcriptomic regionalization of the tibialis anterior (TA) described in Chapter 2, we next investigated whether similar spatial patterns might also be evident at the epigenetic level. Specifically, we hypothesized that the proximal-distal differences in gene expression and metabolic specialization^10^ could, at least in part, be attributable to underlying variation in DNA methylation across these regions. Alternatively, if methylation did not recapitulate the spatial organization observed at the RNA and metabolic levels^10^, it might instead reflect broader regulatory features, such as sex-specific differences^7-9^. To address these possibilities, our study examined how DNA methylation varies across anatomically distinct sites within the TA and assessed whether these patterns align with the transcriptomic differences previously identified^10^.

To address these questions, we performed genome-wide DNA methylation profiling using MeDseq^11^ on selected TA sections derived from the same tissues used for spatial transcriptomics in Chapter 2. This dataset enabled a direct multi-omic comparison of gene expression and methylation within matched anatomical regions. Our results reveal that, unlike the clear regionalization observed at the transcriptomic level, DNA methylation patterns show minimal spatial variation along the proximal-distal axis. Instead, the dominant differences arise between male and female muscles, indicating that sex is the primary determinant of methylation variation in adult TA tissue. Finally, by integrating MeDseq and RNAseq data, we explore how methylation may influence fiber-type plasticity and contribute to sex-specific muscle phenotypes, providing new insights into the regulatory architecture of skeletal muscle.

## Results

### Methylation status at TSS and gene bodies of regionalized genes along the proximal-distal axis of TA muscles

To investigate the role of DNA methylation in whole skeletal muscle regionalization, we analyzed sections from the same muscles used for spatial transcriptomics^10^ (Figure 1a). For the transcriptomic analysis, odd-numbered sections were processed with TOMOseq^12^, whereas for the spatial methylomic analysis, we selected even-numbered sections and analyzed them with MeDseq. MeDseq is a genome-wide DNA methylation sequencing technique based on digestion of DNA with the methylation-dependent restriction enzyme LpnPI, which recognizes methylated CpG sites and cleaves the DNA 16 bp upstream and downstream of the methylated cytosine, generating short (∼32 bp) fragments (Supplementary Figure 1a)^11^. Following adaptor ligation, size selection, and sequencing, only fragments containing a centrally positioned CpG are retained for analysis. Consequently, the presence of sequencing reads at LpnPI recognition sites indicates methylated CpGs, whereas the absence of reads indicates a lack of detectable methylation at those positions. We analyzed the methylome of 12 sections from 3 muscles (Muscle 1, Muscle 2, and Muscle 3, see Methods, Supplementary Fig. 1) and assigned their spatial identity based on the annotation of the corresponding interleaved sections used in the transcriptomic analysis (Figure 1b). Methylation differences were quantified using mean normalized read counts for each group of sections across all muscles (Methods). As a quality check, we confirmed that methylation read counts were highly correlated across all muscle samples, indicating consistent library preparation and sequencing performance (Supplementary Fig. 1d). We also examined coverage along gene bodies and observed the expected depletion of CpG methylation around TSS (Supplementary Fig. 1e).

**Figure 1.**
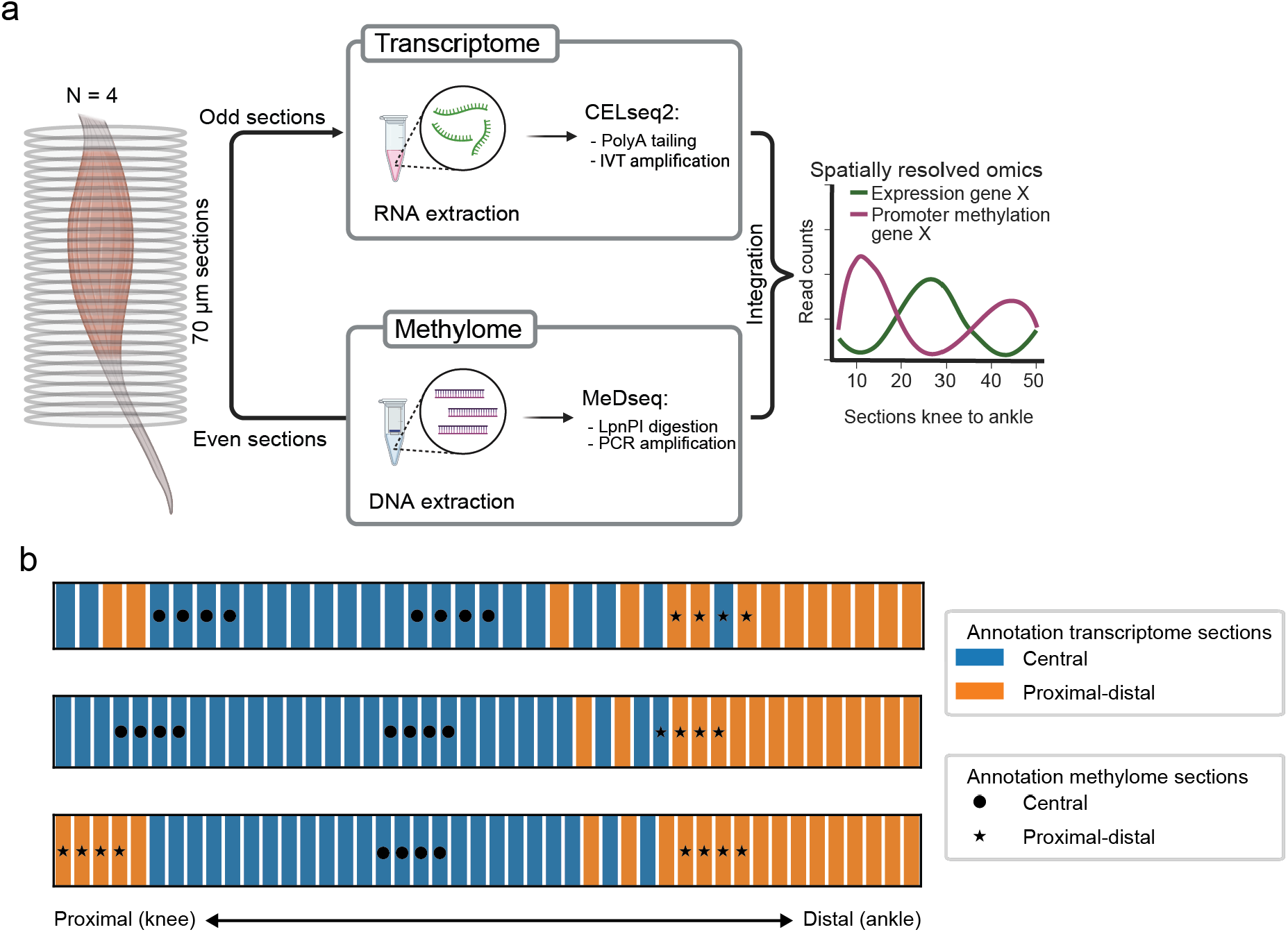
Spatial analysis of transcriptome and methylome from whole TA muscles. a) Schematic overview of the experimental workflow. Tibialis anterior muscles were cryosectioned perpendicular to the proximal-distal axis into 70 μm slices. Odd-numbered sections were used for RNA extraction and spatial transcriptomic analysis, whereas genomic DNA was extracted from even-numbered sections for methylome profiling. b) Sections analyzed by TOMOseq are shown from proximal (knee) to distal (ankle) and colored according to transcriptomic annotation (orange: proximal-distal; blue: central, see Chapter 2 for more information). Selected sections for MeDseq are overlaid on their corresponding preceding sections and annotated based on spatial identity determined from the transcriptomic analysis. Symbols indicate classification of sections: stars denote proximal-distal and circles denote central.

To test whether spatial gene expression patterns are associated with differential DNA methylation, we compared methylation levels between proximal-distal and central sections for genes identified as regionally expressed by TOMOseq, examining CpGs within TSS (3 kb upstream and 100 bp downstream) and gene bodies. Analyses were performed both at the level of individual CpG sites and by pooling all CpGs for each gene in TSS regions or gene bodies. If methylation patterns mirrored spatial gene expression, we would expect methylation at TSS regions to show an inverse trend relative to gene expression, whereas gene body methylation would follow the same trend. However, comparisons of methylation coverage between regions revealed no discernible spatial patterns distinguishing the two regions (Figure 2a,b). Pearson correlation analysis confirmed this lack of spatial differentiation, showing highly similar methylation levels between central and proximal-distal sections (Figure 2c,d) at both individual CpG resolution and when aggregated across TSS regions or gene bodies. Together, these results demonstrate a lack of differences in methylation levels across these differentially expressed genes between proximal-distal and central sections which suggests that DNA methylation does not underlie the spatial regionalization of gene expression along the proximal-distal axis.

**Figure 2.**
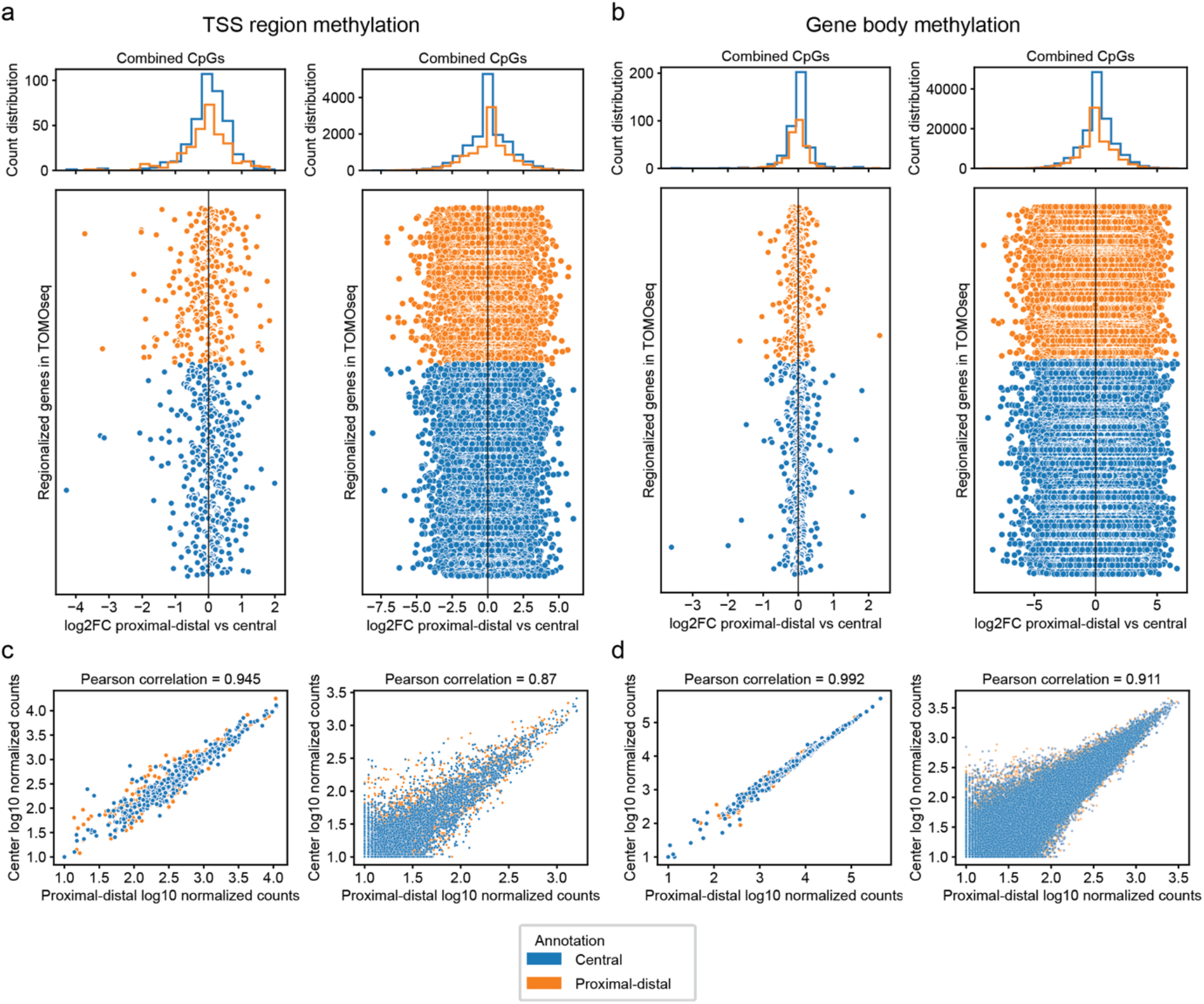
Methylation status of regionalized genes from Chapter 2. a-b) Scatter plots and histograms showing DNA methylation differences between proximal-distal and central regions for each regionalized gene from Chapter 2. The log_2_ fold change (log_2_FC) was calculated from the mean normalized coverage across all annotated proximal-distal or central sections combined from all three muscles. In the scatter plots (bottom), each point represents methylation difference in the TSS region (a) or gene body (b) for a gene, either at individual CpG sites (right) or after merging CpGs within each region (left). Genes up-regulated in central sections are shown in blue, and those up-regulated in proximal-distal sections in orange (see chapter 2 for more information). The order of genes on the y-axis is the same for all scatter plots. The histograms (top) show the distribution of log_2_FC values across all genes for each annotation, providing an overview of the methylation shift between regions. c-d) Pearson correlation of normalized counts for each regionalized gene between central and proximal-distal sections of all three muscles for TSS region (c) and gene body (d) methylation in the sum of all CpG sites from the same gene (left) or the individual CpG sites (right). Colors indicate if the gene was up-regulated in central (blue) or proximal-distal (orange) sections (see chapter 2 for more information).

### Effect of sex in methylation levels of proximal-distal and central sections

Previous studies have reported substantial sex-specific differences in DNA methylation in skeletal muscle^7-9^. We considered the possibility that such sex-dependent variation could obscure potential differences in methylation across proximal-distal and central sections. To address this, we incorporated sex information into our analysis: Muscle 1 corresponded to Female 1, Muscle 2 to Female 2, and Muscle 3 to Male 1.

Pearson correlation analysis across sections from all muscles, independent of spatial annotation or sex, revealed clear differences between male and female samples, but no consistent differences between proximal-distal and central sections within either sex (Figure 3). Notably, male sections exhibited higher correlations than female sections, likely due to the lower coverage in female sections, which reduces the signal-to-noise ratio and, as a result, decreases similarity between female sections. Specifically, methylation in TSS regions did not exhibit spatial patterns that could explain the regionalized gene expression identified by TOMO-seq (Figure 4a,b, Supplementary Fig. 2a,b). Similarly, gene body methylation levels were consistent across central and proximal-distal sections, independent of sex (Figure 4c,d, Supplementary Fig. 2c,d). These results indicate that sex-specific methylation does not mask spatial differences in methylation along the proximal-distal axis, but instead represents the dominant source of methylation variation in adult TA muscle. Together with our previous analyses, this further supports the conclusion that DNA methylation is unlikely to be a primary driver of spatial regionalization of gene expression in adult TA muscles.

**Figure 3.**
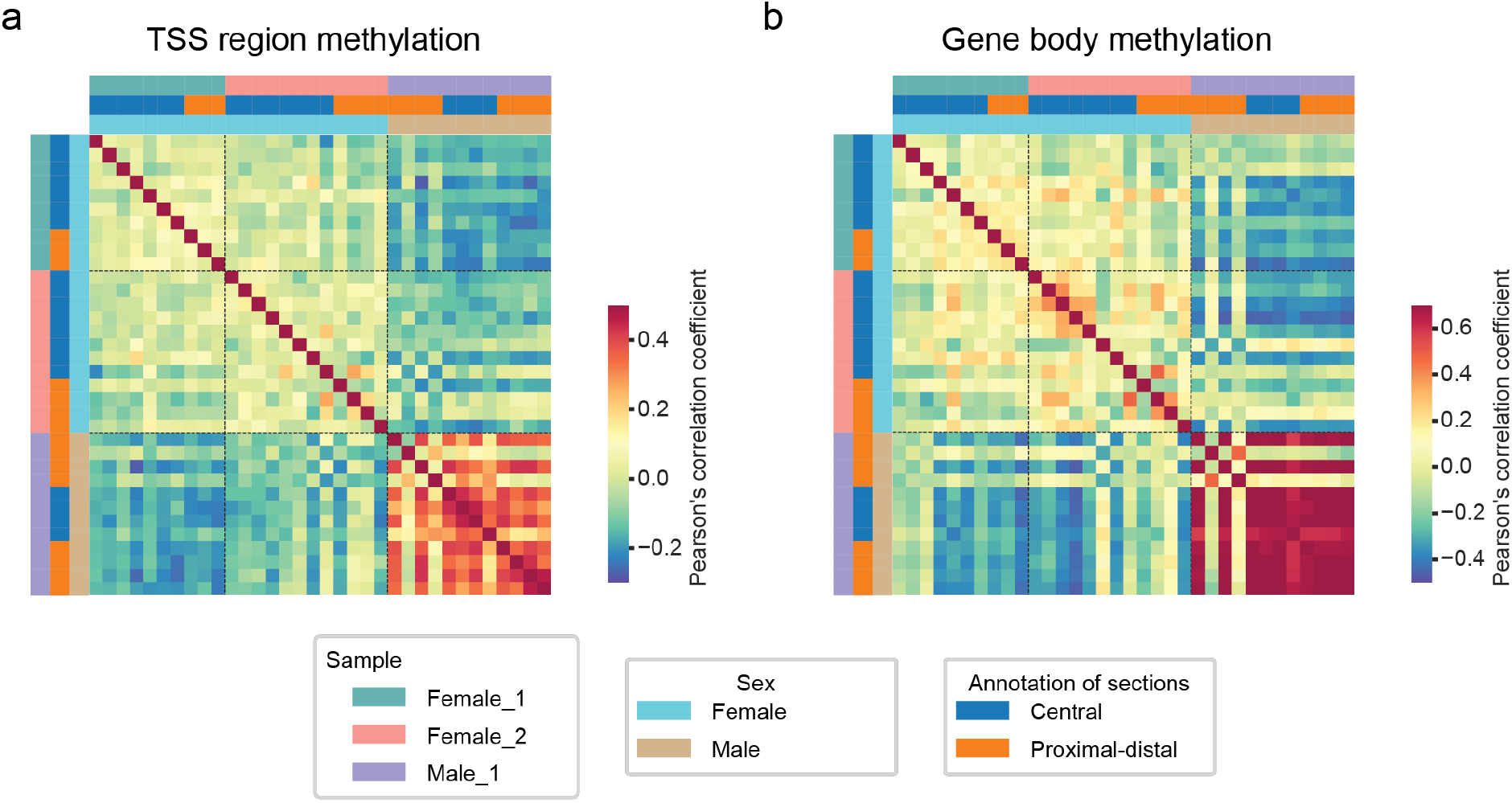
Methylation differences are explained by sex differences. a-b) Person correlation heatmap of all pair-wise comparison of sections for CpG sites in TSS regions (a) or gene bodies (b).

**Figure 4.**
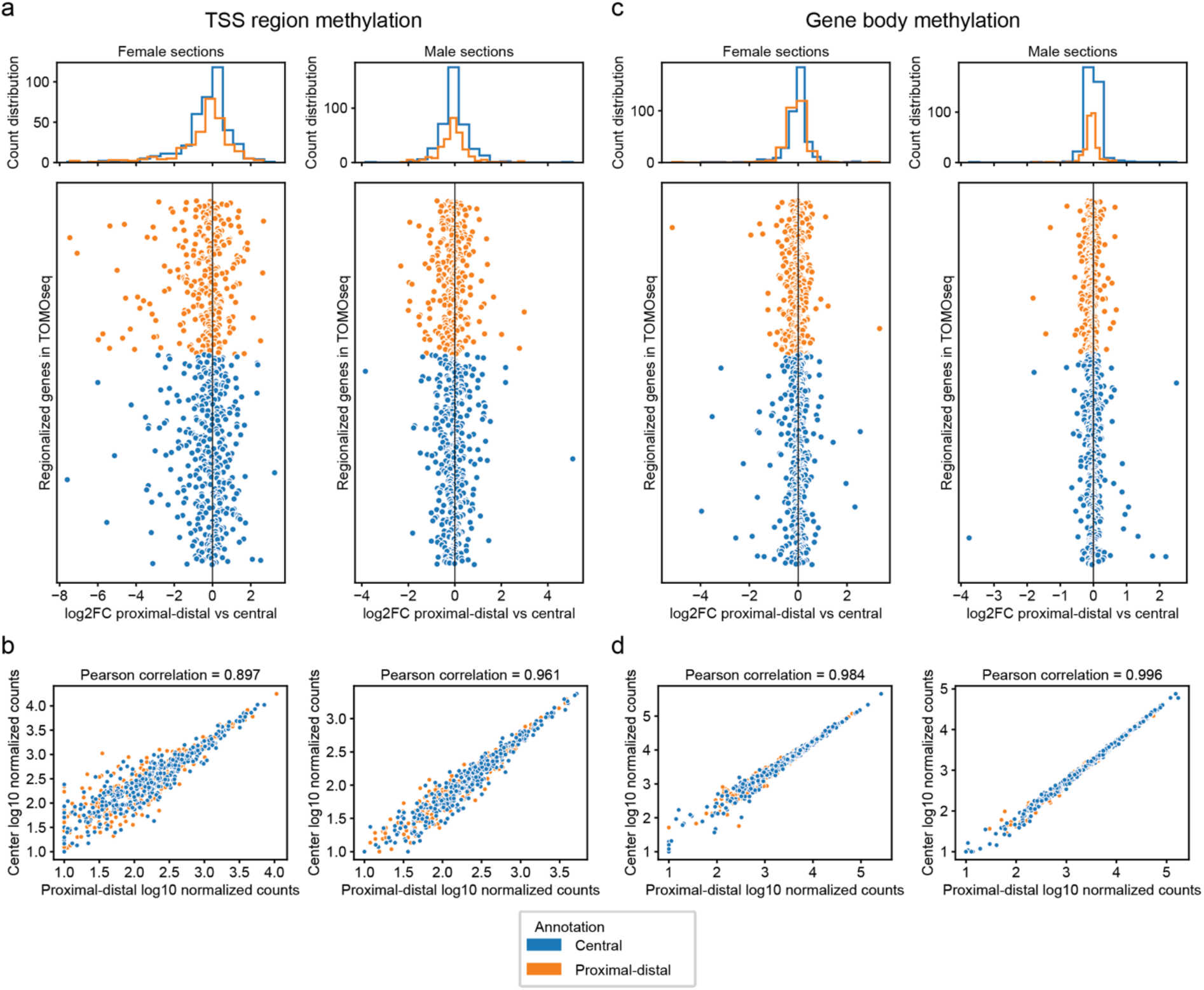
Methylation status of regionalized genes by sex group in combined CpGs. a-c) Scatter plots and histograms showing DNA methylation differences between proximal-distal and central regions for each regionalized gene from Chapter 2 in female (left) or male (right) sections. The log_2_ fold change (log_2_FC) was calculated from the mean normalized coverage across all annotated proximal-distal or central sections for female muscles combined (Muscle 1 and Muscle 2) or for the male muscle (Muscle 3). In the scatter plots (bottom), each point represents methylation difference in the TSS region (a) or gene body (c) for a gene, after merging CpGs within each region. Genes up-regulated in central sections are shown in blue, and those up-regulated in proximal-distal sections in orange (see chapter 2 for more information). The order of genes on the y-axis is the same for all scatter plots. The histograms (top) show the distribution of log_2_FC values across all genes for each annotation, providing an overview of the methylation shift between regions. b-d) Pearson correlation of normalized counts for each regionalized gene between central and proximal-distal sections of female muscles combined (Muscle 1 and Muscle 2, right) or for the male muscle (Muscle 3, left) for TSS region (b) and gene body (d) methylation in the sum of all CpG sites from the same gene. Colors indicate if the gene was up-regulated in central (blue) or proximal-distal (orange) sections (see chapter 2 for more information).

### Methylation differences are associated to sex differences

To investigate the factors that contribute to differences in DNA methylation, we applied a linear model with interaction effects, including sex, spatial location (central vs. proximal-distal annotation), and their interaction. Instead of analyzing individual CpG sites, we quantified DNA methylation in consecutive 1,000 bp genomic segments, providing a consistent resolution across the genome. The analysis was performed separately for segments overlapping with TSS regions, gene bodies, and regulatory regions (promoters, promoter flanking regions, transcription factor binding sites, open chromatin regions, and CTCF binding sites). This approach allowed us to quantify the relative contribution of each factor to methylation variation in different genomic contexts.

Across TSS regions, sex was the primary determinant of methylation differences, with 8,873 bins showing a significant effect, whereas spatial annotation and the interaction term were significant in only 7 and 3 bins, respectively (Figure 5a). Interestingly, 6,934 bins showed a significant overall model fit, even though none of the individual predictors (sex, spatial annotation, or their interaction) reached statistical significance on their own. This pattern may reflect subtle combined effects of the variables that are not attributable to any single factor, but it may also indicate limited statistical power due to the relatively small number of sections analyzed, which could make it difficult to detect smaller effect sizes for individual coefficients. Sex also emerged as the dominant differentiating factor in gene bodies, with 105,770 bins showing significant differences (Figure 5b). Spatial annotation explained methylation in only 9 bins, and the interaction term in 75 bins. Similarly, 90,427 bins showed significance in the overall model without any single coefficient being individually significant. Analysis of regulatory regions revealed a consistent pattern, with 42,360 regions significantly associated with sex, 30 with spatial annotation, and 22 with the interaction term (Figure 5c). In addition, 33,942 bins in regulatory regions were significant in the overall model without individual coefficient significance. Collectively, these results indicate that sex is the predominant driver of variation in DNA methylation in adult TA muscles, whereas spatial position along the proximal-distal axis and its interaction with sex contribute minimally. These findings are consistent with our earlier analyses showing little to no spatial differences in methylation for genes exhibiting region-specific expression.

**Figure 5.**
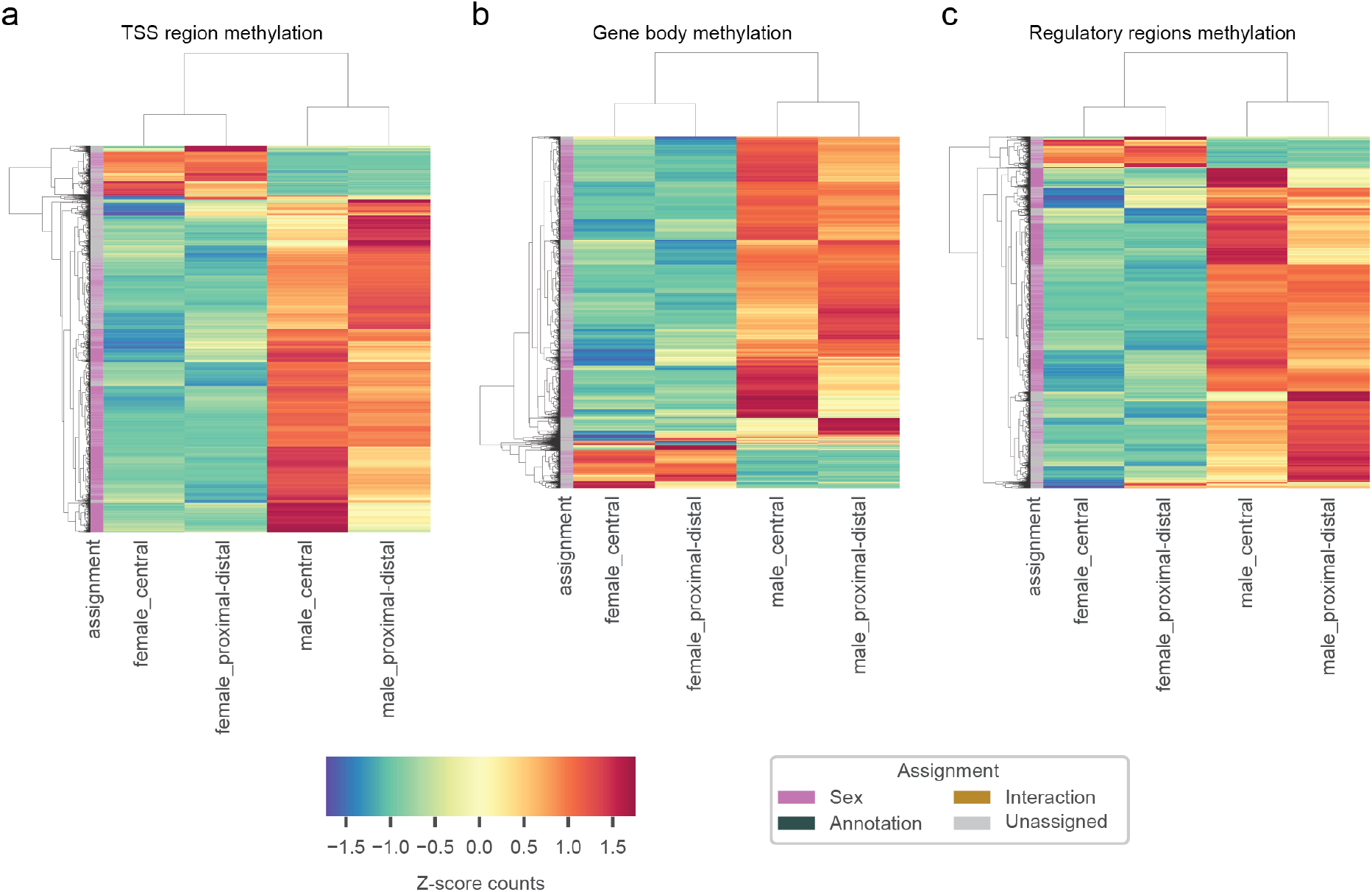
Factors driving DNA methylation differences across TA muscle sections. Heatmaps show the z-score of average methylation read counts in 1,000 bp bins for female_central, male_central, female_proximal-distal, and male_proximal-distal sections. Only regions significant in the overall linear model with interaction effects are included (adjusted *P* value < 0.05). Rows are genomic regions (1000 bp bins), columns are sections, and hierarchical clustering reflects similarity in methylation patterns. Bins are color-coded by significant term: sex, annotation (spatial position), interaction, or unassigned (significant overall but not for individual coefficients). Panels show different genomic contexts: (a) TSS regions, 15,814 bins; (b) gene bodies, 196,206 bins; (c) regulatory regions, 76,328 bins.

To further investigate sex-specific methylation differences, we performed a classical differential methylation analysis comparing male and female sections. For each 1000 bp bin, we computed the average methylation read counts across male or female sections and applied a two-sample t-test. The null hypothesis (H_0_) was that there is no association between sex and methylation levels, whereas the alternative hypothesis (H_1_) assumed a sex-dependent difference. Bins with adjusted *P* value < 0.05 were considered significantly differentially methylated between sexes. We merged significant bins falling within the same gene for TSS regions and gene bodies. Regions were further filtered based on the absolute log_2_ fold-change > 2 (Supplementary Table 1-3). Heatmaps of these significant regions revealed that, across TSS regions, gene bodies, and regulatory regions, a majority of regions exhibited higher methylation in male sections compared to female sections, independent of spatial annotation along the proximal-distal axis (Figure 6). Although we performed GO enrichment analysis on these sex-associated regions, no terms passed statistical significance. We therefore processed to examine whether this sex-biased pattern aligned with known biological differences. As expected, the methylation profiles mirrored established sex differences in fiber-type composition, with male muscles enriched in fast-glycolytic fibers and female muscles enriched in fast-oxidative or slow-oxidative fibers^13^. In particular, the glycolytic marker *Aldoa* showed higher methylation in the TSS region of female sections (male vs female log2FC = -1.03), whereas the slow skeletal muscle myosin light chain isoform *Myl3* had more methylation in male sections (male vs female log2FC = 1.17). For gene body methylation, we observed the opposite trend: male sections exhibited higher methylation at the glycolytic gene *Pkm* (log2FC = 2.15), while the oxidative gene *Idh2* showed higher methylation in female sections (log2FC = –1.47).Together, these patterns highlight sex as a dominant factor influencing DNA methylation in adult TA muscles, while spatial position has a comparatively minor effect.

**Figure 6.**
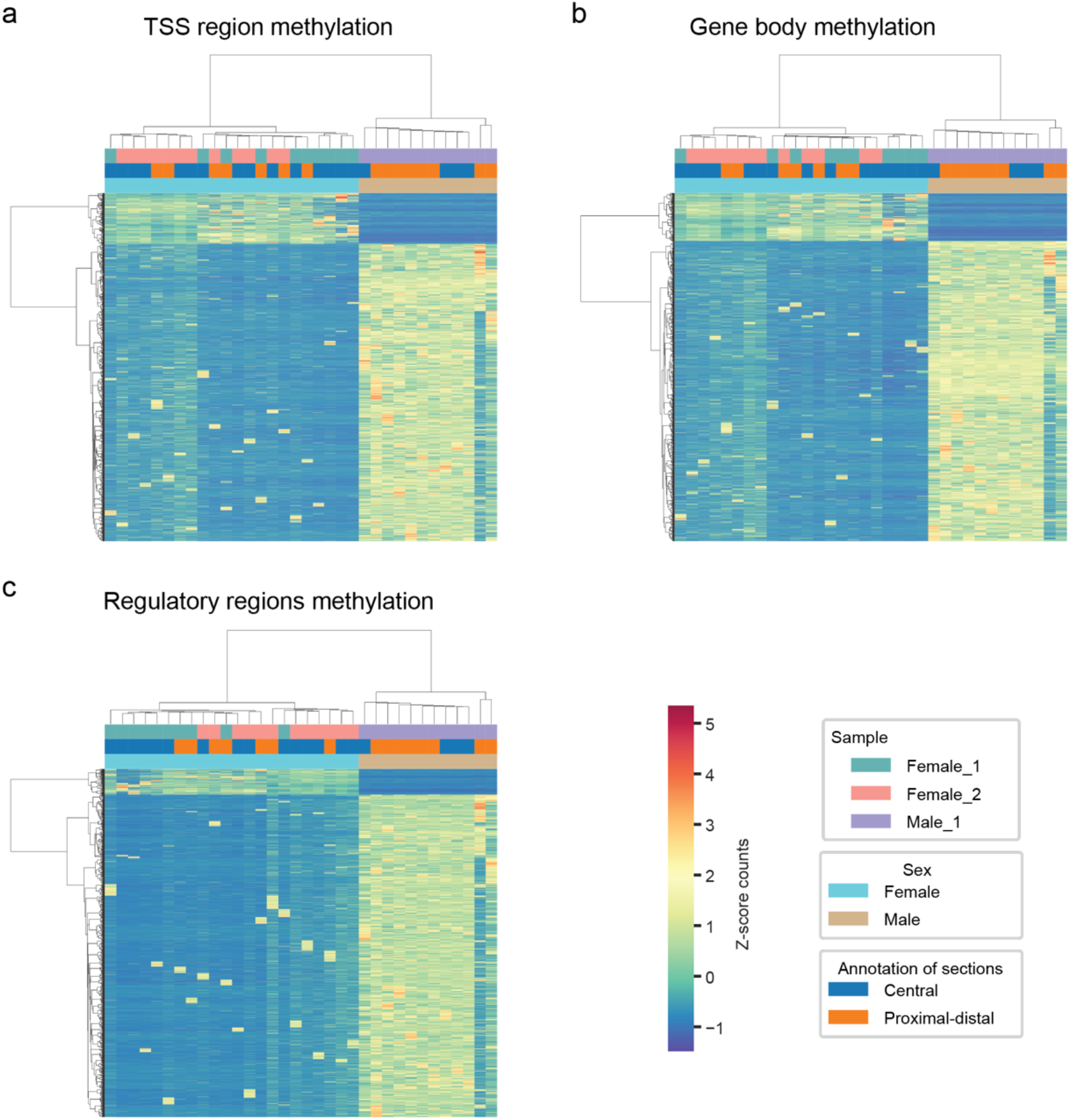
Heatmaps of differentially methylated regions between male and female TA muscle sections. Heatmaps show the z-score of normalized methylation read counts in regions identified as significantly differentially methylated between male and female sections (two-sample t-test adjusted *P* value < 0.05 and absolute log_2_ fold-change > 2). Columns correspond to individual sections, grouped by sample (Muscle 1, Muscle 2 and Muscle 3), sex (male vs. female) and spatial annotation (central vs. proximal-distal). Rows represent genomic 1000 bp bins, and hierarchical clustering was performed to visualize similarities in methylation patterns. Colors indicate z-score normalized methylation levels, highlighting sex-specific differences across regions. Panels display different genomic contexts: (a) TSS regions, 5,792 genes; (b) gene bodies, 6,075 genes; (c) regulatory regions, 35,063 regions.

### Integration of transcriptomic and methylome: Role of DNA methylation in fiber type acquisition

To examine whether sex-specific DNA methylation differences are reflected at the transcriptional level, we performed differential gene expression analysis between muscles within and across sexes. For this analysis, Muscle 4 was included as a second male dataset (Male 2) to enable balanced comparisons, even though corresponding methylation data were not available for this muscle.

Gene expression patterns were highly similar across muscles within the same sex, with only 21 differentially expressed genes (DEGs) detected between the two female muscles and 88 DEGs between the two male muscles (Figure 7a). In contrast, all female-male comparisons revealed substantially larger transcriptional differences (Figure 7b), ranging from 537 to 1,121 DEGs depending on the specific pairwise comparison. Among these sex-associated differences, we detected consistent shifts in fiber-type markers: the oxidative *Myh2* and intermediate *Myh1* isoforms were upregulated in females, whereas the glycolytic marker *Pkm* and *Kdm5d* (Y-linked chromosome gene involved in sex-specific gene expression) were upregulated in males. These patterns agree with previously reported sex differences in muscle fiber and metabolic profiles^13,14^. These results indicate that sex is a major determinant of transcriptional variation in adult TA muscle, and they align with our observations of widespread sex-biased DNA methylation differences across promoters, gene bodies, and regulatory regions.

**Figure 7.**
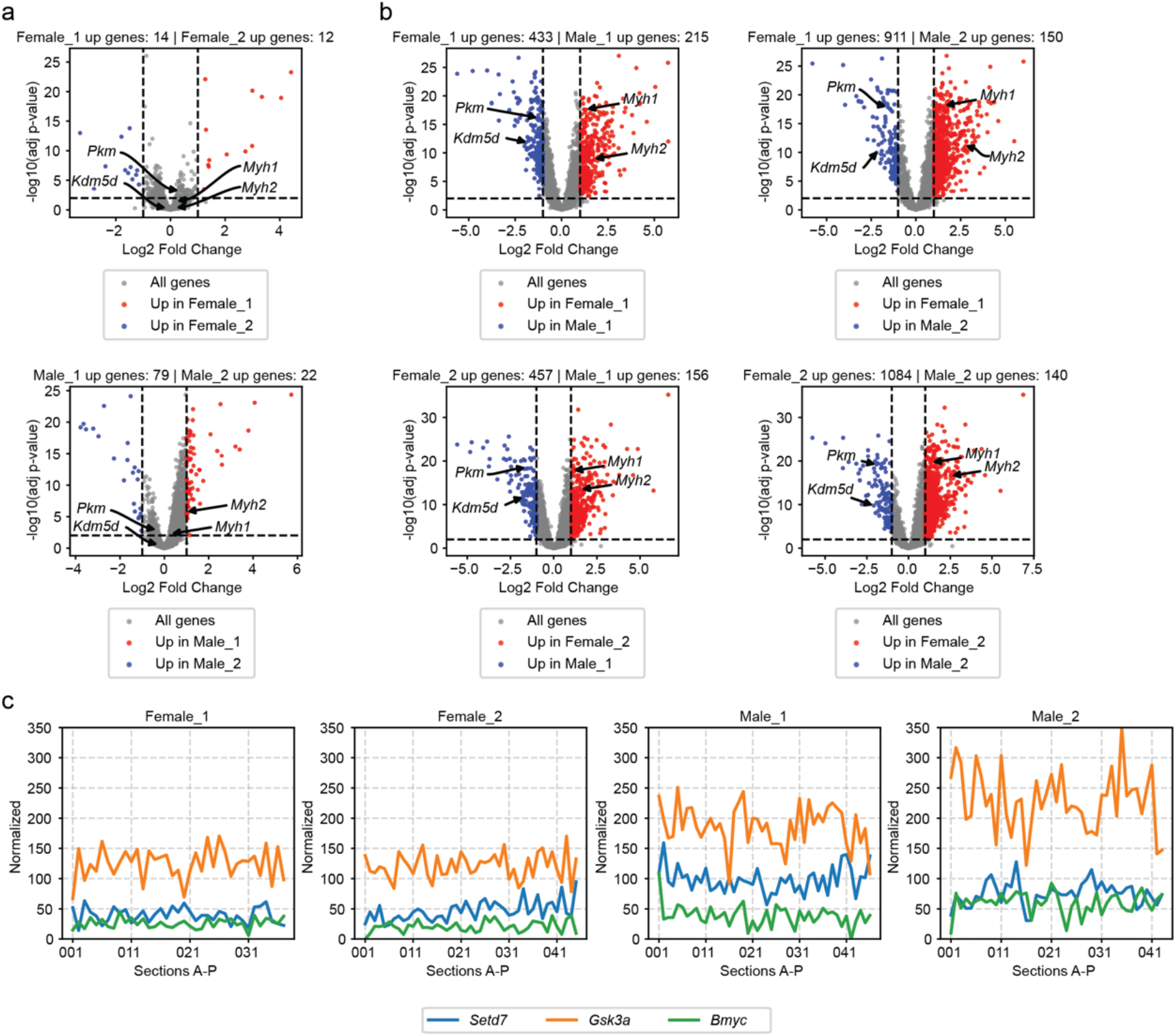
Genes with differential expression between male and females that can explain methylation differences. a) Differential expressed gene (DEG) analysis between muscles of the same sex. b) DEG analysis between all pairwise male and female muscle combinations. c) Normalized expression levels of methylation related genes along the proximal-distal axis of TA muscles.

To explore whether these broad methylation differences could be linked to sex-biased expression of genes with known epigenetic or regulatory functions, we examined DEGs with established roles in chromatin modification, DNA methylation, or muscle fiber identity. Several candidates emerged as biologically compelling (Figure 7c).

First, *Setd7*, a histone-lysine methyltransferase, was consistently upregulated in male muscles (Figure 7c). SETD7 (SET domain-containing protein 7) is a lysine methyltransferase that is known to regulate sarcomere organization and myosin heavy chain-II (MyHC-II) expression (fast-twitch fibers), implicating it directly in fiber-type specification^15^. Given that methylation differences were particularly pronounced in regions associated with fiber-type-specific genes^5^, we suggest a potential correlation between the increased *Setd7* expression in males and the male-biased promoter hypermethylation and fiber-type regulation pattern observed. This might also more generally show a link between transcriptional control of chromatin state and observed methylation patterns.

Second, *Gsk3a* displayed robust male-enriched expression (Figure 7c). *Gsk3a* has been implicated in regulating DNA methylation at imprinted loci, where its loss reduces global methylation levels^16^. Its role in mTORC1 signaling further connects it to muscle growth and metabolic regulation^17^. Consistent with these known functions, male TA sections showed both higher *Gsk3a* expression and globally elevated methylation (Figure 6), supporting the hypothesis that *Gsk3a* activity may help sustain higher methylation levels in male muscle.

Finally, *Bmyc*, a regulator of the transcription factor *Myc*, was also upregulated in males (Figure 7c). *Myc* is a potent driver of epigenetic remodeling and is known to recruit DNA methyltransferases to its target loci^18^. Specifically, *Myc*-overexpression resulted in reduced *Myh2* mRNA levels in murine skeletal muscle^19^. Therefore, increased *Bmyc* expression may amplify *Myc*-mediated methylation programs in males, offering an additional mechanism through which transcriptional differences can reinforce sex-dependent DNA methylation patterns.

Taken together, these analyses identify a set of sex-biased transcriptional regulators (*Setd7, Gsk3a*, and *Bmyc*) whose expression profiles, known molecular functions, and relationships to methylation pathways align with the male-biased DNA methylation observed across promoters, gene bodies, and regulatory elements. These findings suggest that sex-specific differences in transcription factor and chromatin-modifying gene expression may contribute to shaping the distinct fiber type and metabolic landscapes of male and female TA muscles.

## Discussion

In this study, we investigated the contribution of DNA methylation to transcriptional and functional heterogeneity in adult skeletal muscle, with a particular focus on spatial organization along the proximal-distal axis and potential sex-specific regulatory differences. By integrating genome-wide DNA methylation profiling with matched spatial transcriptomic datasets from the TA, we were able to evaluate how epigenetic variation relates to patterns of gene expression and fiber-type identity.

A central finding of our work is the striking dissociation between spatially regionalized gene expression and underlying DNA methylation patterns. Although TOMOseq analysis in Chapter 2 revealed robust proximal-distal variation in oxidative vs glycolytic metabolism, contractile gene programs, and fiber-type composition^10^, MeDseq profiling showed that CpG methylation remains largely stable across these spatial domains. Neither promoter nor gene-body methylation exhibited the expected inverse or concordant trends that would typically accompany region-specific changes in gene transcription. Correlation analyses and coverage comparisons consistently demonstrated that proximal-distal and central sections were nearly indistinguishable at the methylome level, even when examining CpGs associated with the most strongly regionalized transcripts. These observations indicate that, unlike processes such as developmental patterning or long-term cellular identity, spatial specialization within adult muscle fibers is not driven by static differences in CpG methylation.

Instead, results from a linear regression model incorporating sex, spatial location, and their interaction point to sex as the predominant source of methylation variability in the adult TA muscle. Across TSS regions, gene bodies, and regulatory elements, sex accounted for orders of magnitude more differentially methylated regions than spatial location or sex-by-location interactions. This strikingly sex-biased methylation landscape mirrors several previous genome-wide studies reporting extensive differences between male and female skeletal muscle methylomes^7-9^. Importantly, these methylation differences were not subtle: in most genomic contexts, male samples showed higher methylation levels than female samples, suggesting a broad shift in epigenetic state rather than isolated locus-specific effects. Additionally, the higher methylation in males likely contributed to the greater read recovery in male sections, which in turn may explain the higher similarity observed among male sections compared to female sections (Figure 3).

The transcriptional analyses reinforce this conclusion. While muscles from the same sex displayed highly similar gene expression profiles, comparisons between males and females revealed hundreds of differentially expressed genes, including markers of fiber-type identity and metabolic specialization. Consistent with known physiological differences^13,14^, females showed increased expression of oxidative and intermediate MyHC isoforms (*Myh2* and *Myh1*), whereas males upregulated glycolytic markers and Y-linked transcripts. These findings align with a large body of literature documenting sex differences in substrate utilization, fatigue resistance, and myofiber composition.

An intriguing aspect of our study is the identification of sex-biased regulatory genes that may link transcriptional and epigenetic variation. *Setd7, Gsk3a*, and *Bmyc* were found to be more highly expressed in male muscles and have established roles in chromatin modification, DNA methylation maintenance, and transcription factor recruitment^15,16,18^. Their functions offer biologically plausible mechanisms through which male-biased transcriptional programs could reinforce male-biased methylation patterns. For example, *Gsk3a* has been shown to influence global methylation levels^16^, while *Bmyc* is a well-known modulator of *Myc*, a factor capable of recruiting DNA methyltransferases and reshaping epigenetic landscapes^18^. *Setd7’s* role in fiber-type specification further connects methylation-associated regulatory pathways to functional muscle phenotypes^15^. Although our dataset cannot establish causality, the convergence of gene expression, methylation status, and known molecular functions suggests a coordinated regulatory axis linking sex-specific chromatin states to muscle identity.

Together, these findings advance our understanding of the regulatory architecture of adult skeletal muscle in several important ways. First, they clarify that spatial regionalization along the proximal-distal axis is shaped primarily by transcriptional and metabolic programs rather than by stable CpG methylation differences. This suggests that other regulatory modalities, such as histone modifications, 3D chromatin interactions, local innervation patterns, or regional differences in mechanical or metabolic load, are more likely drivers of anatomical specialization within a single muscle. Second, our results highlight sex as a major determinant of the skeletal muscle methylome, emphasizing that sex must be considered when interpreting epigenetic data. Finally, the integration of methylomic and transcriptomic evidence reveals potential molecular links between sex-specific epigenetic regulation and fiber-type specification, suggesting that chromatin-modifying enzymes and transcriptional regulators may help reinforce the distinct metabolic and contractile properties of male and female muscle.

Several constraints should be considered when interpreting these findings. First, although the gene expression regionalization observed in homeostatic TA muscle could not be explained by differences in TSS region, gene-body, or regulatory methylation, even after accounting for sex, our methylome profiling relied on MeDseq, which captures a substantial portion of CpG sites but does not provide full genome-wide coverage^11^. It is therefore possible that spatially informative CpGs outside the detectable fraction were not assessed. Second, the number of biological replicates for the methylation analysis was limited, with only one male and two female muscles included. While the sex-biased methylation patterns observed were strong and consistent with previous reports, this sample size restricts our ability to fully characterize inter-individual variability or identify subtler sex-dependent effects. Additional replicates, especially from both sexes, will be critical for validating and refining the conclusions drawn here.

Overall, this work underscores the importance of multi-omic approaches for uncovering the mechanisms that shape muscle phenotype. While DNA methylation does not appear to drive the spatial heterogeneity observed within the TA, it plays a prominent role in sex-dependent muscle biology. Future studies leveraging single-nucleus multi-omics, longitudinal training models, or hormonal manipulation will be essential to disentangle how sex hormones, chromatin regulators, and methylation pathways interact to support muscle function across the lifespan.

## Methods

### Animals

DCM-Polr2b:m2rtTA mice^20^ were housed at the animal facility of Erasmus University Medical Center Rotterdam. Pax7nGFP mice^21^ (kindly provided by Dr. S. Tajbakhsh, Pasteur Institute Paris, France) were housed at the animal facility of Leiden University Medical Center. 4 to 6-month-old mice used in this research mice were sacrificed by cervical dislocation. The experiments were evaluated and approved by the local animal welfare committee under DEC numbers 11600, AVD11600202013788 and AVD10100202115681.

### MeDseq

DNA was extracted from selected cryosections using the QIAamp DNA Micro Kit (Qiagen) according to the manufacturer’s protocol for laser-microdissected tissues. MeDseq analyses were performed as previously described^11,20^. At least 10 ng of isolated DNA was used for LpnPI (New England Biolabs) digestion. Stem-loop adapters were blunt-end ligated to the repaired DNA, followed by amplification with a high-fidelity polymerase to incorporate dual-indexed barcodes and generate indexed Illumina next-generation sequencing (NGS) libraries. Amplified products were purified using a Pippin HT system with 3% agarose gel cassettes (Sage Science). Multiplexed libraries were sequenced on an Illumina HiSeq 2500 system to generate single-end 50 bp reads according to the manufacturer’s instructions. Dual-indexed reads were demultiplexed using bcl2fastq (Illumina).

### Pre-processing MeDseq data

Raw FASTQ files were subjected to Illumina adaptor trimming, and reads were filtered based on LpnPI restriction site occurrence between 14 and 19 bp for CpG methylation from either 5′ or 3′ end of the read. Reads were filtered based on the presence of CpG methylation sites (C^me^CG, ^me^CGG, and G^me^CGC, Supplementary Figure 1a). Reads that passed the LpnPI filter were mapped to GRCm38 using BWA. Mapped reads were used to assign read count scores to each individual LpnPI site in the GRCm38 genome and highly methylated CpG sites were excluded based on coverage levels (Supplementary Figure 1b). We excluded sections with low coverage based on the total counts (Supplementary Figure 1c). Genome-wide individual LpnPI site scores were used to generate read count scores for the following annotated regions: transcription start site (TSS, 3 kb before and 100 bp after), gene bodies (from TSS until transcription end site, TES) and regulatory regions (promoters, promoter flanking regions, transcription factor binding sites, open chromatin, CTCF biding sites). Data was binned in non-overlapping 1000 bp windows. For each genomic bin, methylated read counts from all CpG sites within the bin were summed to obtain a bin-level methylation signal. This sum was used for downstream analyses, including differential methylation testing and linear modeling. In some analyses, counts were normalized across sections to library size and scaled to mean coverage of all sections (Supplementary Figure 1c) before comparison of sections.

### Methylation status of regionalized genes

CpG sites located within promoter regions or gene bodies of regionalized genes identified in Chapter 2 were selected for further analysis. Unbinned normalized methylation data were used to calculate mean coverage across sections annotated as proximal-distal or central according to the spatial transcriptomic analysis from the three muscles. For each gene, log_2_ fold-changes (log_2_FC) were computed for proximal-distal versus central mean normalized counts. Pearson correlation coefficients were calculated for each gene to assess the concordance of methylation counts between proximal-distal and central regions. CpG sites were analyzed individually or aggregated by gene to evaluate differential methylation patterns across regions and between sexes. Sex-specific analyses were performed by stratifying samples into male and female groups.

### Linear regression model with interaction

To investigate the contribution of sex and spatial location to DNA methylation, we applied linear regression models to binned methylation data (non-overlapping 1000 bp bins). For each genomic bin, methylation levels were modeled as a function of sex (male vs. female), spatial location (annotation: central vs. proximal-distal), and their interaction effect for each bin:

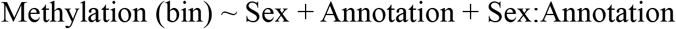

We considered the following regression model:

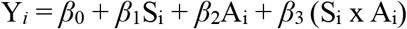

where *Y*_*i*_ is the methylation coverage for section *i*; S_i_ is the sex state (male or female) of section i; A_i_ is the section annotation (central or proximal-distal) of section i; *β*_1_, *β*_2_, and *β*_3_ are the sex, annotation, and interaction effect sizes, respectively; and *ε*_i_ is the residual error term, assumed to be normally distributed with mean 0 and constant variance.

For each bin, regression coefficients and *P* values were extracted for the main effects of sex and spatial location, as well as for the interaction term. Overall model significance was evaluated using the F-test. *P* values were corrected across genes using the Benjamini-Hochberg false discovery rate procedure. Bins were initially selected for further consideration if the overall model was statistically significant at adjusted *P* value < 0.05 (F-test). Among these selected bins, individual contributions of sex, spatial location, and their interaction effects were evaluated using the adjusted *P* values of the individual regression coefficients. Bins were classified according to the factor(s) driving the effect:

- Sex-associated bins: bins where the coefficient for sex had adjusted *P* value < 0.05.
- Spatial location-associated bins: bins where the coefficient for annotation had adjusted *P* value < 0.05.
- Interaction-associated bins: bins where the interaction term coefficient had adjusted *P* value < 0.05.

This strategy allowed quantification of the relative contribution of each factor to methylation variation in different genomic contexts and enabled grouping of bins based on the primary driver of methylation differences.

### Differentially methylated regions

Binned DNA methylation data were pre-processed by separating samples according to sex (male or female). For each genomic bin, the mean methylation levels were compared between groups using a two-sided Welch’s t-test and log_2_ fold-change (log_2_FC) were computed. The null hypothesis (H_0_) assumed no association between sex and methylation level, while the alternative hypothesis (H_1_) assumed a significant difference. T-tests were performed on all bins with at least one non-zero methylation value in either group. *P* values were corrected for multiple testing using the Benjamini-Hochberg procedure to control the false discovery rate. Bins with corrected *P* values < 0.05 and absolute log_2_FC > 2 were considered differentially methylated.

### Differentially expressed genes

Gene expression matrices from individual muscles were first normalized to library size by dividing raw counts by the total number of reads per sample. For each pairwise muscle comparison, genes absent (count = 0) in both muscles were excluded from the analysis. For each gene, log_2_FC was computed adding a pseudocount of 1 to avoid taking the logarithm of zero. Differential expression between muscles was assessed using Welch’s t-test, implemented across sample replicates for each muscle. *P* values were corrected for multiple testing using the Benjamini-Hochberg false discovery rate procedure. Genes were considered differentially expressed if they met both criteria: adjusted *P* value < 0.01, and absolute log_2_FC > 2.

## Acknowledgements

The authors thank the LUMC animal facility and the LUMC Light and Electron Microscopy. Pax7nGFP mice were kindly provided by S. Tajbakhsh (Institut Pasteur, Paris, France).

## Author contributions

N.G. and F.S. conceived and designed the project. C.M.M. carried out the experiments with guidance from F.S. and A.A. throughout. C.M.M. analyzed and interpreted the results with input from F.S., A.A. and N.G. C.M.M. prepared the material for the MeDseq analysis, the experiment was performed by R.B. under the guidance of J.G.. C.M.M. wrote the manuscript and assembled the data with assistance from F.S., A.A. and N.G.

## Competing interests

The authors declare that they have no competing interests

## Funding

This study was supported by The novo Nordisk Foundation Center for Stem Cell Medicine (reNEW) grant NFF21CC0073729.

## Supplementary material

**Supplementary Figure 1.**
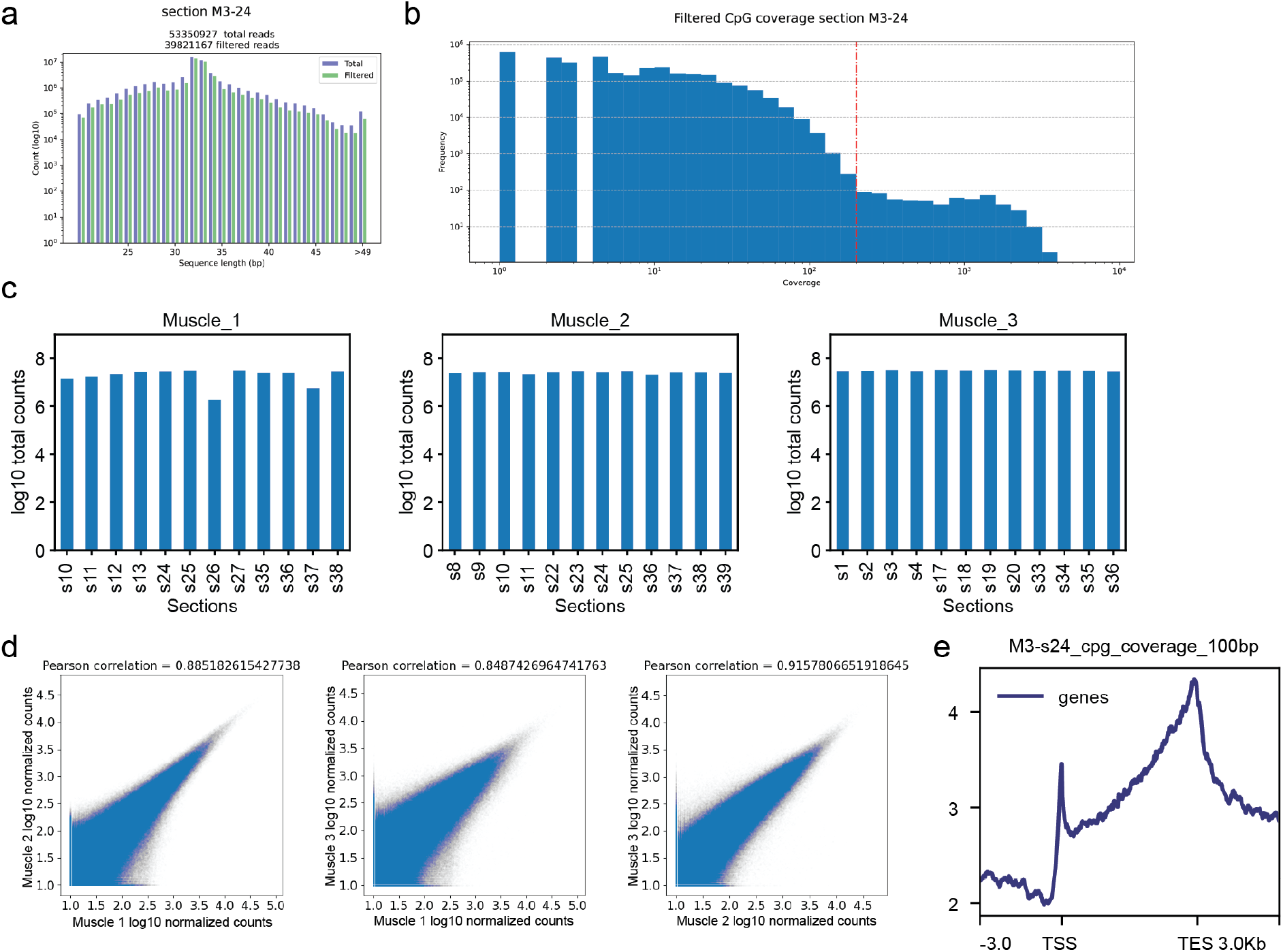
MeDseq data pre-processing. a) Bar plot of the read count before (total) and after trimming and LpnPI filtering for one representative section. b) Representative coverage plot of one section to select threshold for filtering CpG sites with high coverage. c) Total coverage in each section. d) Pearson correlation of total counts for each pair-wise comparison between muscle samples. e) Methylation read counts were aggregated for all annotated genes, spanning from 5 kb upstream of the transcription start site (TSS) to 5 kb downstream of the transcription end site (TES). The line represents mean coverage across genes.

**Supplementary Figure 2.**
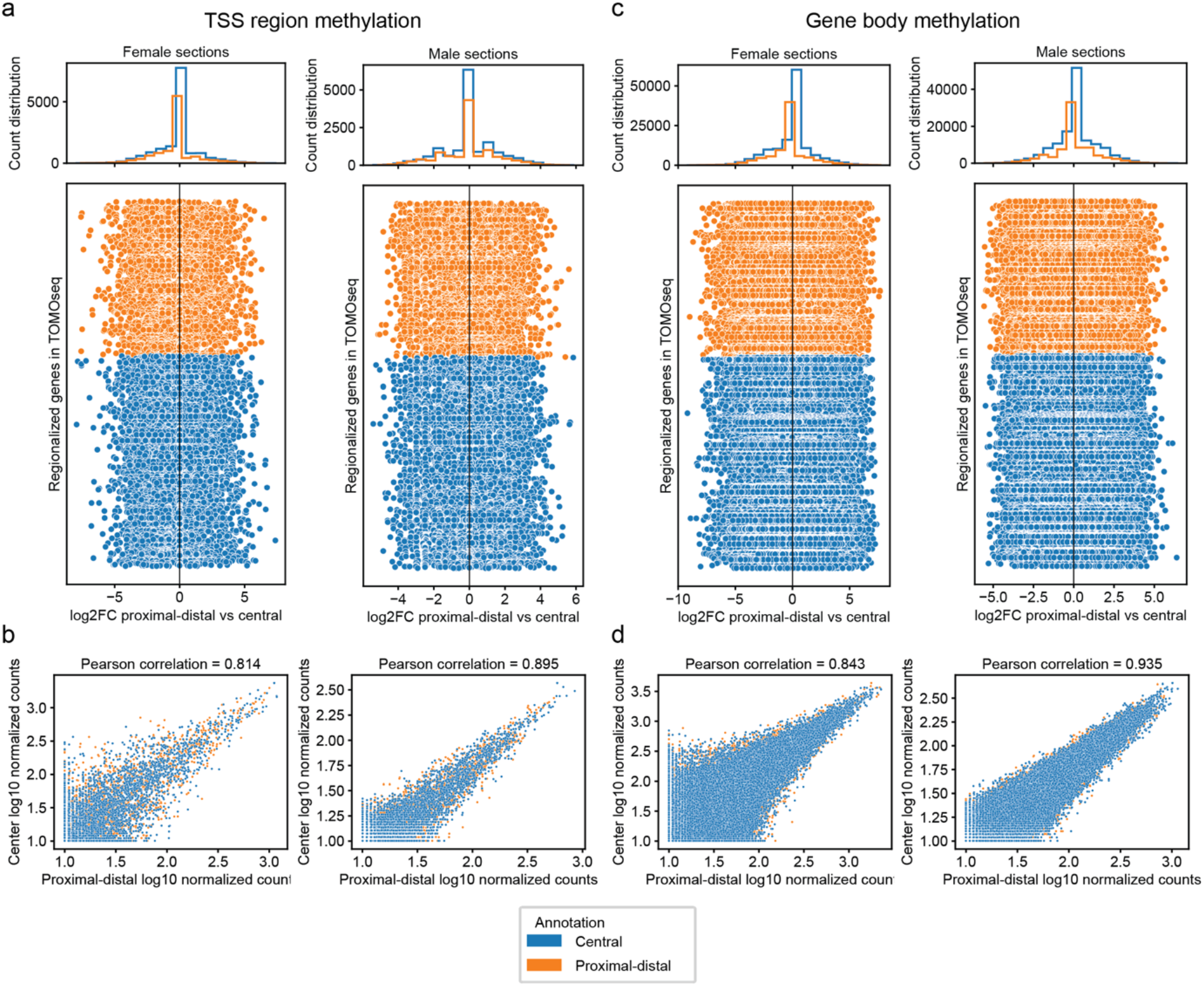
Methylation status of regionalized genes by sex group in individual CpGs. a-c) Scatter plots and histograms showing DNA methylation differences between proximal-distal and central regions for each regionalized gene from Chapter 2 in female (left) or male (right) sections in individual CpG sites. The log_2_ fold change (log_2_FC) was calculated from the mean normalized coverage across all annotated proximal-distal or central sections for female muscles combined (Muscle 1 and Muscle 2) or for the male muscle (Muscle 3). In the scatter plots (bottom), each point represents methylation difference in the TSS region (a) or gene body (c) for each CpG sites of one gene. Genes up-regulated in central sections are shown in blue, and those up-regulated in proximal-distal sections in orange (see chapter 2 for more information). The order of genes on the y-axis is the same for all scatter plots. The histograms (top) show the distribution of log_2_FC values across all genes for each annotation, providing an overview of the methylation shift between regions. b-d) Pearson correlation of normalized counts for each regionalized gene between central and proximal-distal sections of female muscles combined (Muscle 1 and Muscle 2, right) or for the male muscle (Muscle 3, left) for TSS region (b) and gene body (d) methylation in each CpG site from the same gene. Colors indicate if the gene was up-regulated in central (blue) or proximal-distal (orange) sections (see chapter 2 for more information).

**Supplementary Table 1.**
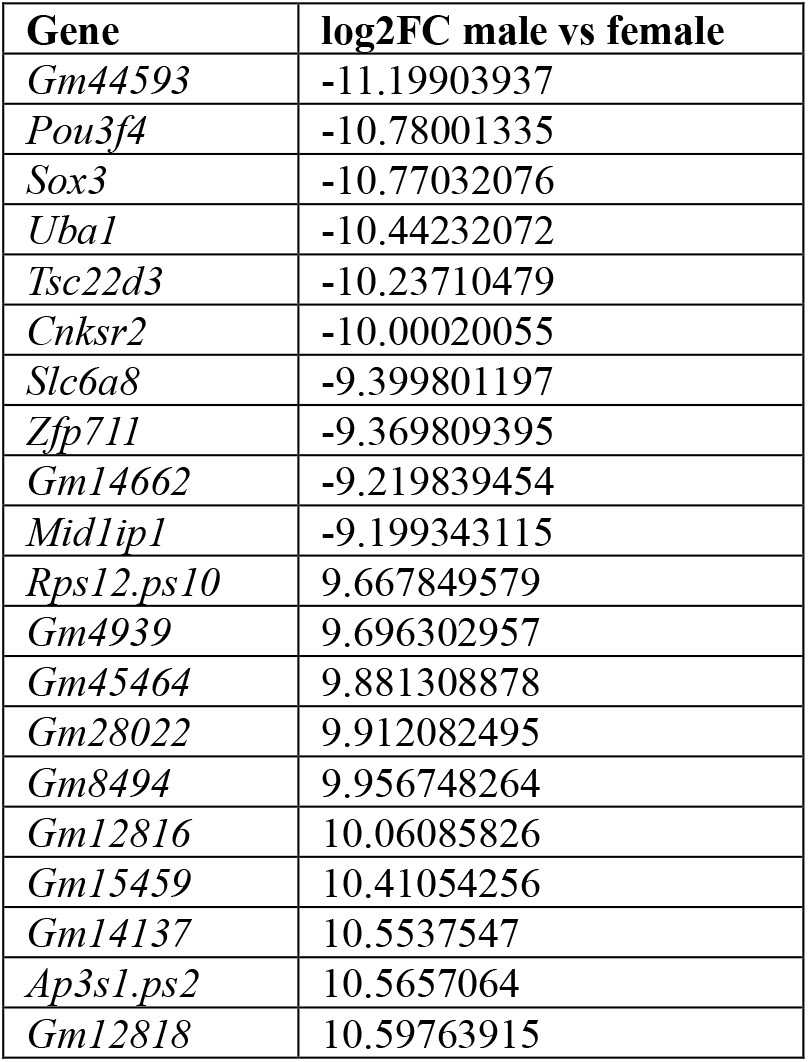
Top 20 DMR in TSS regions between male and female sections.

**Supplementary Table 2.**
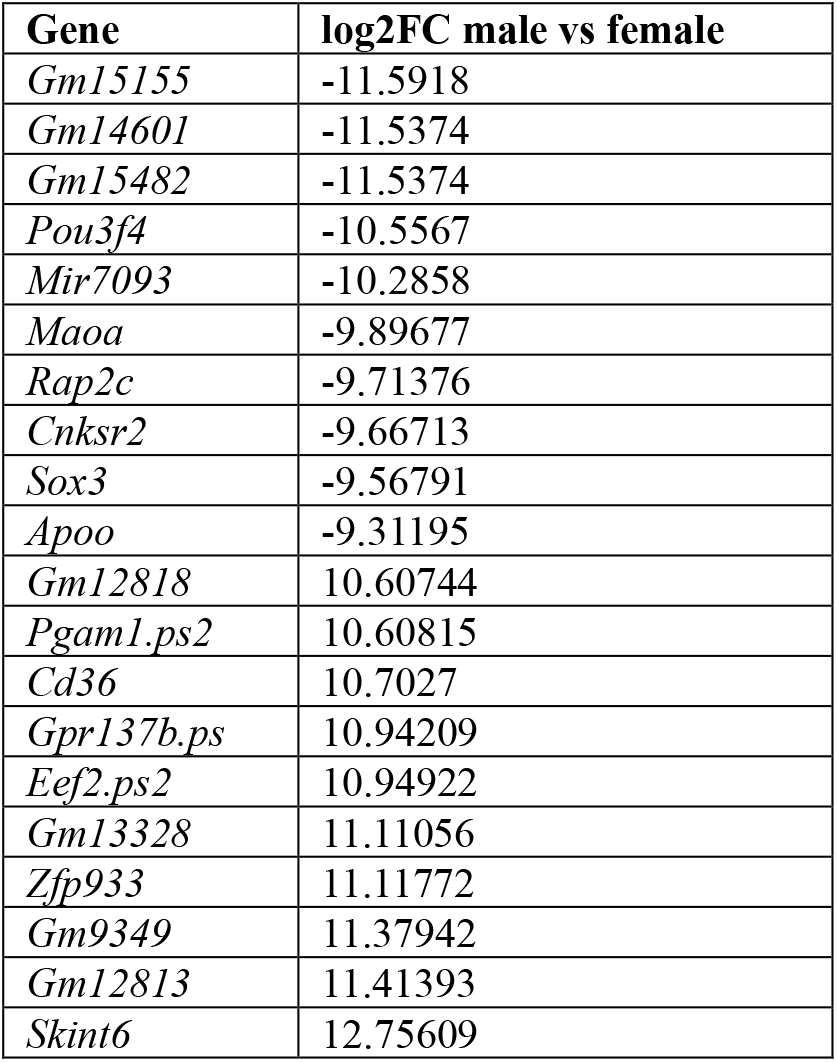
Top 20 DMR in gene body regions between male and female sections.

**Supplementary Table 3.**
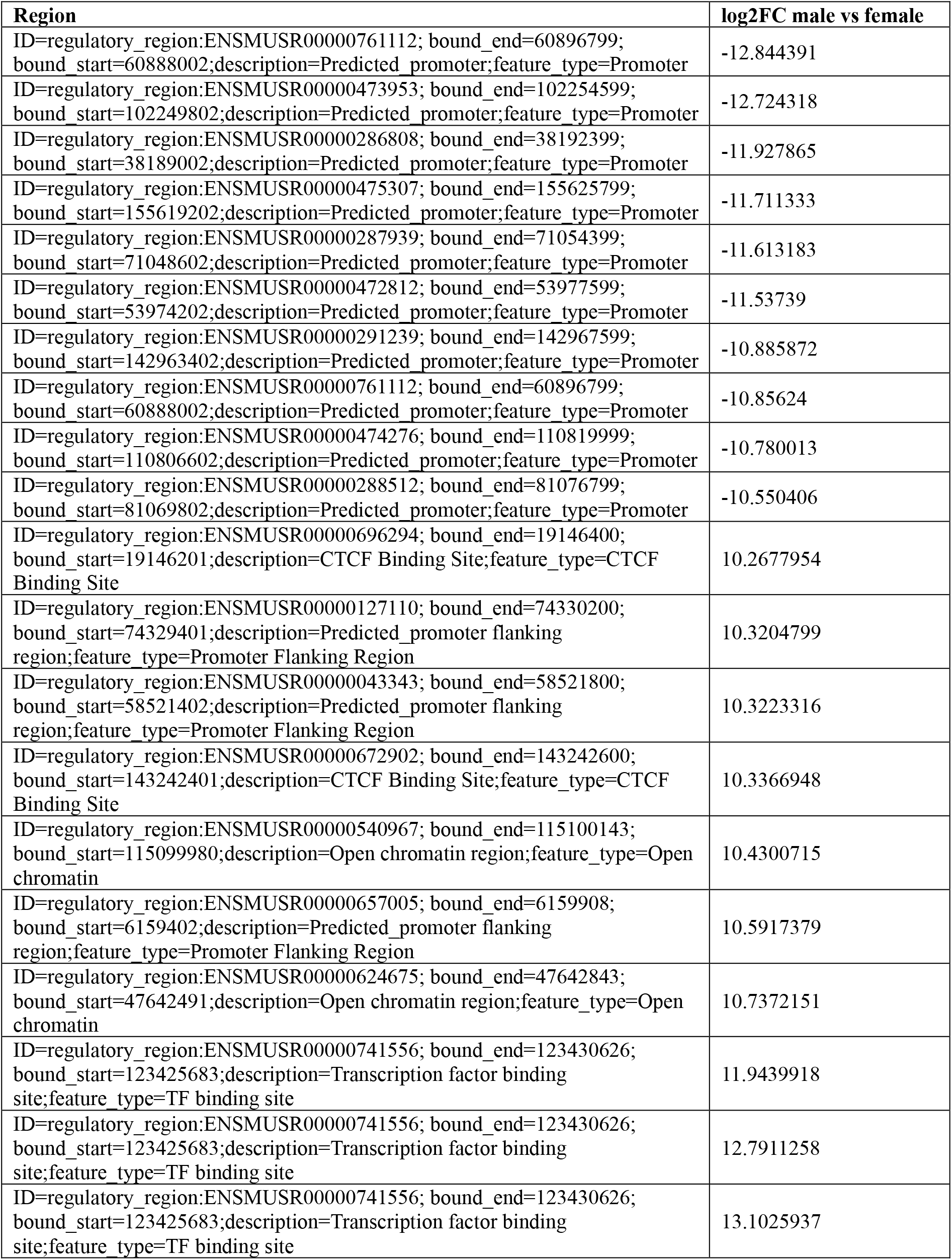
Top 20 DMR in regulatory regions between male and female sections.

## Notes

### Competing Interest Statement

The authors have declared no competing interest.

### Summary of Updates

In the originally published version of this article, the name of the author was incorrectly written as Joost Gribnay. The correct spelling is Joost Gribnau. This has now been corrected in the online version of the article.

## References

1. Mattei, A. L., Bailly, N. & Meissner, A. DNA methylation: a historical perspective. Trends Genet 38, 676–707 (2022).

2. Wang, Q. et al. Gene body methylation in cancer: molecular mechanisms and clinical applications. Clin Epigenetics 14, 154 (2022).

3. Jones, P. A. Functions of DNA methylation: islands, start sites, gene bodies and beyond. Nat Rev Genet 13, 484–492 (2012).

4. Iio, H. et al. DNA maintenance methylation enzyme Dnmt1 in satellite cells is essential for muscle regeneration. Biochem Biophys Res Commun 534, 79–85 (2021).

5. Begue, G., Raue, U., Jemiolo, B. & Trappe, S. DNA methylation assessment from human slow- and fast-twitch skeletal muscle fibers. J Appl Physiol (1985) 122, 952–967 (2017).

6. Geiger, C. et al. DNA methylation of exercise-responsive genes differs between trained and untrained men. BMC Biol 22, 147 (2024).

7. Landen, S. et al. Skeletal muscle methylome and transcriptome integration reveals profound sex differences related to muscle function and substrate metabolism. Clin Epigenetics 13, 202 (2021).

8. Davegardh, C. et al. Sex influences DNA methylation and gene expression in human skeletal muscle myoblasts and myotubes. Stem Cell Res Ther 10, 26 (2019).

9. Landen, S. et al. Sex differences in muscle protein expression and DNA methylation in response to exercise training. Biol Sex Differ 14, 56 (2023).

10. Martinez Mir, C. et al. Spatial multi-omics in whole skeletal muscle reveals complex tissue architecture. Commun Biol 7, 1272 (2024).

11. Boers, R. et al. Genome-wide DNA methylation profiling using the methylation-dependent restriction enzyme LpnPI. Genome Res 28, 88–99 (2018).

12. Junker, J. P. et al. Genome-wide RNA tomography in the zebrafish embryo. Cell 159, 662–675 (2014).

13. Haizlip, K. M., Harrison, B. C. & Leinwand, L. A. Sex-based differences in skeletal muscle kinetics and fiber-type composition. Physiology (Bethesda) 30, 30–39 (2015).

14. O’Reilly, J. et al. Sex differences in skeletal muscle revealed through fiber type, capillarity, and transcriptomics profiling in mice. Physiol Rep 9, e15031 (2021).

15. Nayak, A. et al. Regulation of SETD7 Methyltransferase by SENP3 Is Crucial for Sarcomere Organization and Cachexia. Cell Rep 27, 2725–2736 e2724 (2019).

16. Meredith, G. D. et al. Glycogen synthase kinase-3 (Gsk-3) plays a fundamental role in maintaining DNA methylation at imprinted loci in mouse embryonic stem cells. Mol Biol Cell 26, 2139–2150 (2015).

17. Zhou, J. et al. GSK-3alpha is a central regulator of age-related pathologies in mice. J Clin Invest 123, 1821–1832 (2013).

18. Brenner, C. et al. Myc represses transcription through recruitment of DNA methyltransferase corepressor. EMBO J 24, 336–346 (2005).

19. Murach, K. A. et al. Multi-transcriptome analysis following an acute skeletal muscle growth stimulus yields tools for discerning global and MYC regulatory networks. J Biol Chem 298, 102515 (2022).

20. Boers, R. et al. Retrospective analysis of enhancer activity and transcriptome history. Nature Biotechnology 41, 1582–1592 (2023).

21. Sambasivan, R. et al. Distinct regulatory cascades govern extraocular and pharyngeal arch muscle progenitor cell fates. Developmental Cell 16, 810–821 (2009).

