## Supplementary materials for "Sex-specific DNA methylation in adult skeletal muscle"

### Supplementary material

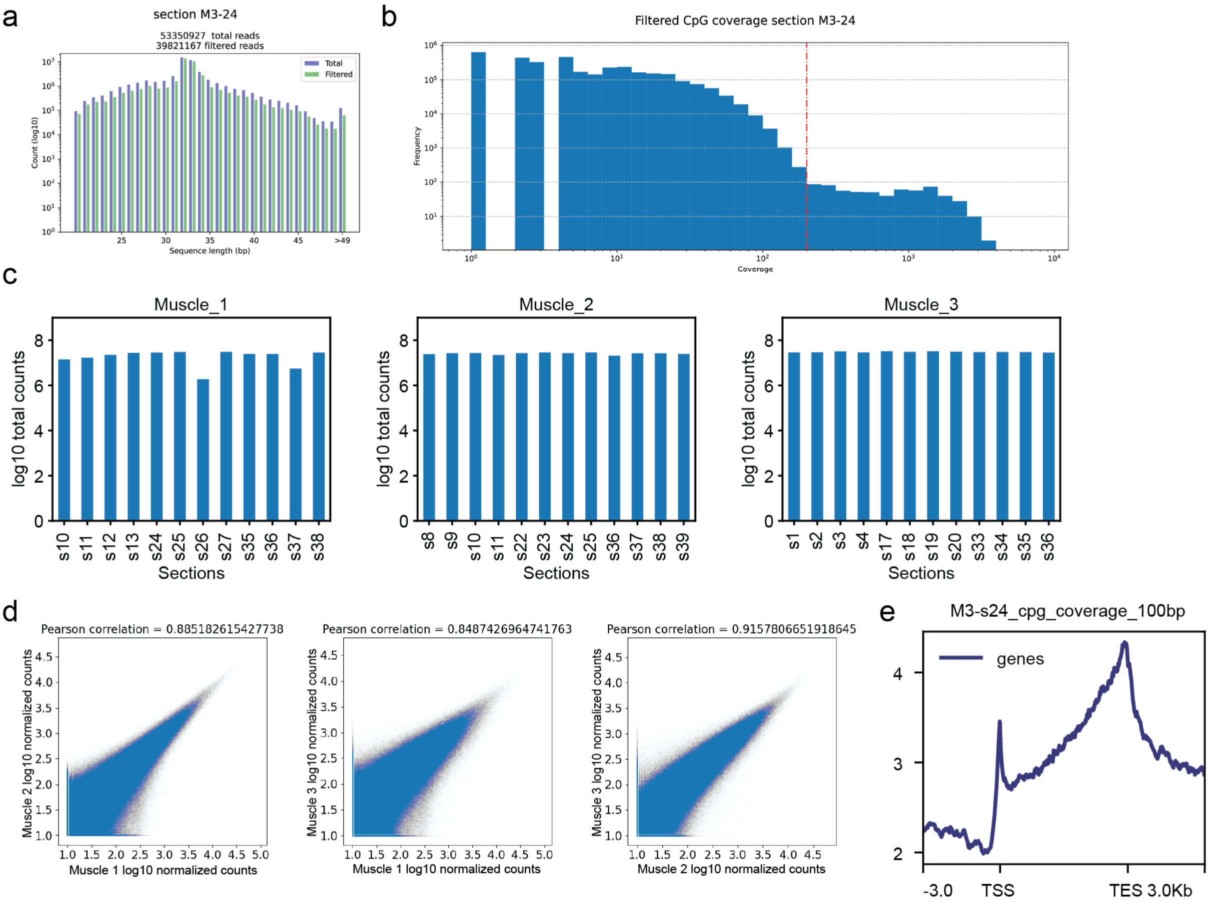


**Supplementary Figure 1. MeDseq data pre-processing.** a) Bar plot of the read count before (total) and after trimming and LpnPI filtering for one representative section. b) Representative coverage plot of one section to select threshold for filtering CpG sites with high coverage. c) Total coverage in each section. d) Pearson correlation of total counts for each pair-wise comparison between muscle samples. e) Methylation read counts were aggregated for all annotated genes, spanning from 5 kb upstream of the transcription start site (TSS) to 5 kb downstream of the transcription end site (TES). The line represents mean coverage across genes.

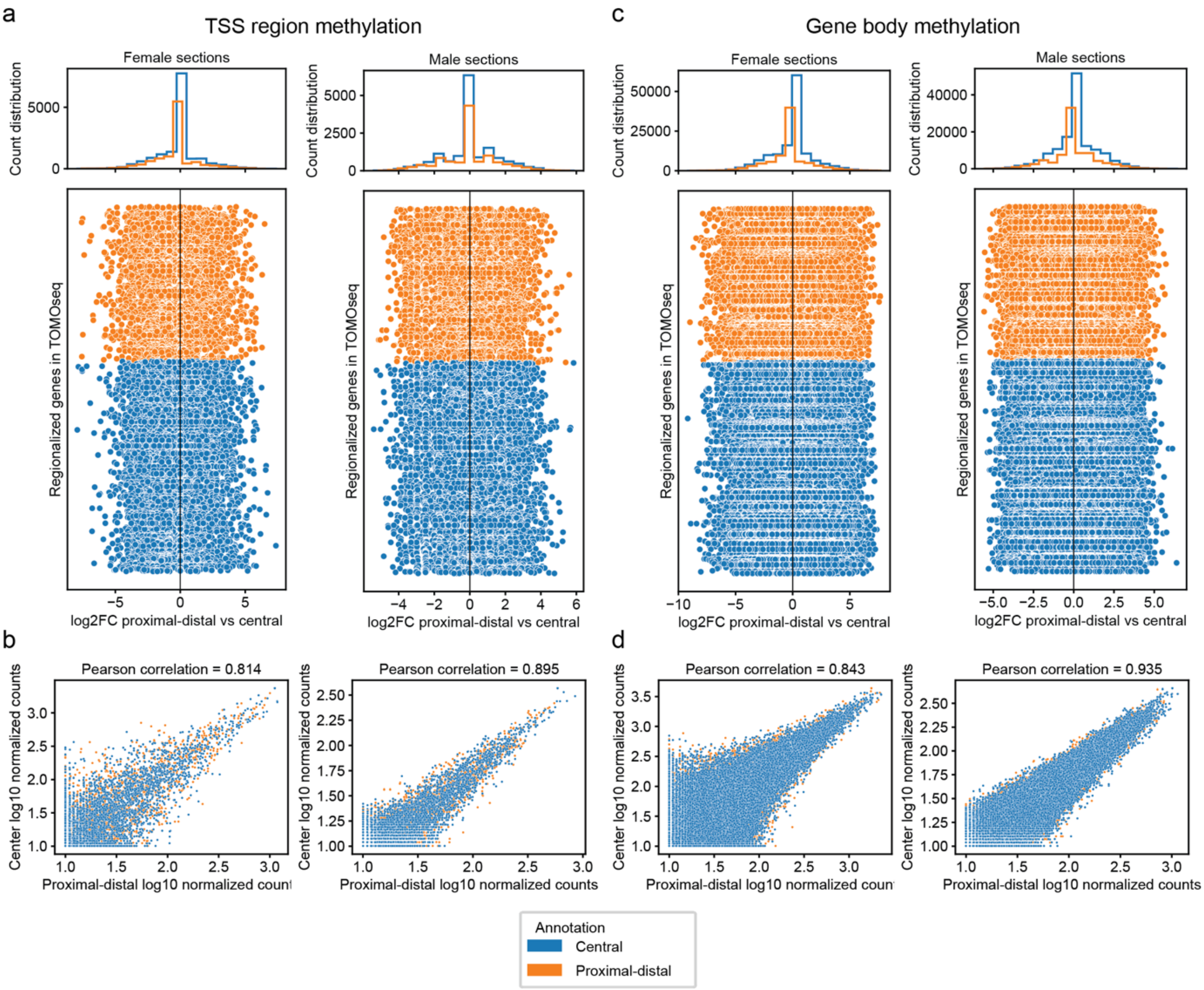


**Supplementary Figure 2. Methylation status of regionalized genes by sex group in individual CpGs.** a-c) Scatter plots and histograms showing DNA methylation differences between proximal-distal and central regions for each regionalized gene from Chapter 2 in female (left) or male (right) sections in individual CpG sites. The $\log_2$ fold change ($\log_2$FC) was calculated from the mean normalized coverage across all annotated proximal-distal or central sections for female muscles combined (Muscle 1 and Muscle 2) or for the male muscle (Muscle 3). In the scatter plots (bottom), each point represents methylation difference in the TSS region (a) or gene body (c) for each CpG sites of one gene. Genes up-regulated in central sections are shown in blue, and those up-regulated in proximal-distal sections in orange (see chapter 2 for more information). The order of genes on the y-axis is the same for all scatter plots. The histograms (top) show the distribution of $\log_2$FC values across all genes for each annotation, providing an overview of the methylation shift between regions. b-d) Pearson correlation of normalized counts for each regionalized gene between central and proximal-distal sections of female muscles combined (Muscle 1 and Muscle 2, right) or for the male muscle (Muscle 3, left) for TSS region (b) and gene body (d) methylation in each CpG site from the same gene. Colors indicate if the gene was up-regulated in central (blue) or proximal-distal (orange) sections (see chapter 2 for more information).

**Supplementary Table 1. Top 20 DMR in TSS regions between male and female sections.**

| Gene | log2FC male vs female |
|---|---|
| *Gm44593* | -11.19903937 |
| *Pou3f4* | -10.78001335 |
| *Sox3* | -10.77032076 |
| *Uba1* | -10.44232072 |
| *Tsc22d3* | -10.23710479 |
| *Cnksr2* | -10.00020055 |
| *Slc6a8* | -9.399801197 |
| *Zfp711* | -9.369809395 |
| *Gm14662* | -9.219839454 |
| *Mid1ip1* | -9.199343115 |
| *Rps12.ps10* | 9.667849579 |
| *Gm4939* | 9.696302957 |
| *Gm45464* | 9.881308878 |
| *Gm28022* | 9.912082495 |
| *Gm8494* | 9.956748264 |
| *Gm12816* | 10.06085826 |
| *Gm15459* | 10.41054256 |
| *Gm14137* | 10.5537547 |
| *Ap3s1.ps2* | 10.5657064 |
| *Gm12818* | 10.59763915 |

**Supplementary Table 2. Top 20 DMR in gene body regions between male and female sections.**

| Gene | log2FC male vs female |
|---|---|
| *Gm15155* | -11.5918 |
| *Gm14601* | -11.5374 |
| *Gm15482* | -11.5374 |
| *Pou3f4* | -10.5567 |
| *Mir7093* | -10.2858 |
| *Maoa* | -9.89677 |
| *Rap2c* | -9.71376 |
| *Cnksr2* | -9.66713 |
| *Sox3* | -9.56791 |
| *Apoo* | -9.31195 |
| *Gm12818* | 10.60744 |
| *Pgam1.ps2* | 10.60815 |
| *Cd36* | 10.7027 |
| *Gpr137b.ps* | 10.94209 |
| *Eef2.ps2* | 10.94922 |
| *Gm13328* | 11.11056 |
| *Zfp933* | 11.11772 |
| *Gm9349* | 11.37942 |
| *Gm12813* | 11.41393 |
| *Skint6* | 12.75609 |

**Supplementary Table 3. Top 20 DMR in regulatory regions between male and female sections.**

| Region | log2FC male vs female |
|---|---|
| ID=regulatory_region:ENSMUSR00000761112; bound_end=60896799; bound_start=60888002;description=Predicted_promoter;feature_type=Promoter | -12.844391 |
| ID=regulatory_region:ENSMUSR00000473953; bound_end=102254599; bound_start=102249802;description=Predicted_promoter;feature_type=Promoter | -12.724318 |
| ID=regulatory_region:ENSMUSR00000286808; bound_end=38192399; bound_start=38189002;description=Predicted_promoter;feature_type=Promoter | -11.927865 |
| ID=regulatory_region:ENSMUSR00000475307; bound_end=155625799; bound_start=155619202;description=Predicted_promoter;feature_type=Promoter | -11.711333 |
| ID=regulatory_region:ENSMUSR00000287939; bound_end=71054399; bound_start=71048602;description=Predicted_promoter;feature_type=Promoter | -11.613183 |
| ID=regulatory_region:ENSMUSR00000472812; bound_end=53977599; bound_start=53974202;description=Predicted_promoter;feature_type=Promoter | -11.53739 |
| ID=regulatory_region:ENSMUSR00000291239; bound_end=142967599; bound_start=142963402;description=Predicted_promoter;feature_type=Promoter | -10.885872 |
| ID=regulatory_region:ENSMUSR00000761112; bound_end=60896799; bound_start=60888002;description=Predicted_promoter;feature_type=Promoter | -10.85624 |
| ID=regulatory_region:ENSMUSR00000474276; bound_end=110819999; bound_start=110806602;description=Predicted_promoter;feature_type=Promoter | -10.780013 |
| ID=regulatory_region:ENSMUSR00000288512; bound_end=81076799; bound_start=81069802;description=Predicted_promoter;feature_type=Promoter | -10.550406 |
| ID=regulatory_region:ENSMUSR00000696294; bound_end=19146400; bound_start=19146201;description=CTCF Binding Site;feature_type=CTCF Binding Site | 10.2677954 |
| ID=regulatory_region:ENSMUSR00000127110; bound_end=74330200; bound_start=74329401;description=Predicted_promoter flanking region;feature_type=Promoter Flanking Region | 10.3204799 |
| ID=regulatory_region:ENSMUSR00000043343; bound_end=58521800; bound_start=58521402;description=Predicted_promoter flanking region;feature_type=Promoter Flanking Region | 10.3223316 |
| ID=regulatory_region:ENSMUSR00000672902; bound_end=143242600; bound_start=143242401;description=CTCF Binding Site;feature_type=CTCF Binding Site | 10.3366948 |
| ID=regulatory_region:ENSMUSR00000540967; bound_end=115100143; bound_start=115099980;description=Open chromatin region;feature_type=Open chromatin | 10.4300715 |
| ID=regulatory_region:ENSMUSR00000657005; bound_end=6159908; bound_start=6159402;description=Predicted_promoter flanking region;feature_type=Promoter Flanking Region | 10.5917379 |
| ID=regulatory_region:ENSMUSR00000624675; bound_end=47642843; bound_start=47642491;description=Open chromatin region;feature_type=Open chromatin | 10.7372151 |
| ID=regulatory_region:ENSMUSR00000741556; bound_end=123430626; bound_start=123425683;description=Transcription factor binding site;feature_type=TF binding site | 11.9439918 |
| ID=regulatory_region:ENSMUSR00000741556; bound_end=123430626; bound_start=123425683;description=Transcription factor binding site;feature_type=TF binding site | 12.7911258 |
| ID=regulatory_region:ENSMUSR00000741556; bound_end=123430626; bound_start=123425683;description=Transcription factor binding site;feature_type=TF binding site | 13.1025937 |
